# Application of DNA barcodes and spatial analysis in conservation genetics and modeling of Iranian Salicornia genetic resources

**DOI:** 10.1101/2020.10.12.335935

**Authors:** Mehrshad Zeinalabedini, Nayer Azam Khoshkholgh Sima, Mohammad Reza Ghaffari, Ali Ebadi, Maryam Farsi

## Abstract

Iran is one of the origins of some Salicornia species. Nevertheless, so far, comprehensive research has not been conducted on genetic potential, distribution, selection of populations, and the economic utilization of Salicornia in Iran. In the current study, Salicornia was collected based on the previous research locations available on 26 different geographical locations of provinces in Iran. Subsequently, an accurate model was provided for identifying of Salicornia species biodiversity by performing spatial analysis and modeling of distributed areas. The results provided valuable information on the diversity of specific geographical regions, conservation status of existing species, prioritization of conservation areas, and selection of areas for Agro-Ecological, leading to the development of industrial agriculture. Further, we validated genes in the field of DNA barcoding in Salicornia plants using matK, rbcL, trnH-psbA, ycf and ITS2 identifying species groups. Together, integrating our results will provide useful information for the management and utilization of Salicornia genetic resources in Iran.

## Introduction

The world’s population is 7 billion in 2018, is expected to rise by 35%, and reaches to 9 billion in 2050. Food and Agriculture Organization of United Nations (FAO) estimates crop production must increase by at least 60% in the future [1] to provide the food and energy demand for the growing population. However, the use of genotypes with high-yield and resistance to biotic and abiotic stress has gradually replaced indigenous varieties and local genotypes endangering access and exploitation of resource and food production for the future [2]. Thus, low genetic diversity may reduce the opportunity to identify and use new sources responding to future challenges, including new pests and pathogens, as well as climate changes [3]. Today, soil salinity has increased due to the unconventional cultivation and irrigation. Moreover, the level of arable land increased dramatically from 8 million to more than 220 million hectares, of which about 45 million hectares are salt-affected to varying degrees [4,5]. The highest saline soils in Asia are in China, India, Pakistan, and Iran [6]. In Iran, salinity is a limiting factor of sustainable agricultural production, covering 34 percent of arid and semi-arid regions of the country, especially in the central, south, and plain of Khuzestan in varying degrees of salinity [7]. Furthermore, about 20%-50% of the arable land is also affected by salinity [8,9]. Therefore, the improvement of agricultural and horticultural crops to increase salinity tolerance and the use of salt-tolerant plants are needed to increase food production, raise employment, and support sustainable development [10]. Halophytes, which include more than 600 taxa of the various genus and species [11] complete their life cycle under high salt concentration (at least 200 mM NaCl) [12,13] making them a rich resource of new crops for the future. Among the common halophytes, *Salicornia* sp. is relatively important due to the high potential for domestication and cultivation [14]. Salicornia is a succulent halophyte, which grows naturally on mangrove swamps as well as seashores. It has also evaluated as a vegetable, forage, and oilseed crop in the agronomic field trials [15]. *Salicornia* sp. is widespread across Iran, mainly in central, western, and northern parts. There is a morphological variation across the genus distribution range of *Salicornia* sp. Some economically feasible applications have been suggested for Salicornia species. They also have suitable biomass for human and domestics consumption [16,17]. Furthermore, Salicornia seed contains 26%–33% oil, of similar composition to safflower oil, along with 30%–33% protein [18]. Therefore, identification and selection of native Salicornia species might be a viable strategy for agricultural development against Iran’s climate change. The number of Salicornia species is between 25 to 30 species worldwide [19]. Hence, the taxonomy classification of Salicornia is very complicated as there is not any general published morphological descriptor for all accepted species. Thus, it often makes it impossible for a non-expert. Previously, some morphological characteristics like growth form, inflorescence, branching of main and lateral stems, lateral flowers and their conditions relative to the main flowers, fruit formation, flowering, and fruiting characteristics as well as dry biomass were used to identify some species of Salicornia with high seed and biomass yield [20–22]. However, a low number of morphological characteristics, phenotypic flexibility, breeding, and hybridization systems are amongst the factors that make Salicornia difficult to distinguish precisely in an area [23]. Currently, few studies were conducted on taxonomic and phylogenetic analysis of the Salicornia genus. The commercial importance and lack of description makes Salicornia a priority for the circumscription of species, subtypes, ecotypes, and natural hybridization in order to develop it as new crop [8]. Teege et al. [8] studied the genetic relationships of intra and inter variation of the two taxa of *Salicornia prompumbens* and *Salicornia stricta* species using AFLP markers [20]. Further, the phylogenetic relationships between eight taxa belonging to the Salicornia genus were examined using the ITS markers [8]. Likewise, the phylogenetic and geographic relationship in different germplasms of Salicornia was evaluated using 6 EST markers [8]. Moreover, genetic diversity of some populations of Salicornia was investigated using RAPD markers [24]. All obtained results showed that these molecular markers have some limitations/disadvantages in accurate taxonomic classifications. Recently, DNA sequencing technology has facilitated the identification of identical genotype with different morphotypes or different genotypes with the same morphotypes [25]. DNA barcoding, which is a new and efficient tool based on the conserved DNA regions of organisms, has provided an efficient and precise method for species-level identifications [26]. DNA barcodes were developed by Consortium for the Barcode of Life (CBOL) as a global standard for the identification of biological species. To date, no comprehensive research has been done on the utility of universal DNA barcodes for all proposed gene regions (either alone or in combination) in Salicornia. The overall aim of our study was to employ three coding plastid regions (*rbcL*, *matK* and *ycf*), one non-coding plastid intergenic spacer regions (*trnH-psbA*), and the internal transcribed spacer of nuclear encoded ribosomal DNA (ITS2) to examine species and population diversity of Iranian Salicornia germplasms. Then, an integrated model was developed to assess Salicornia distribution and best habitat based on climate factors, estimate critical parameters in land management and biodiversity conservation. We assume that our results will open up a new window for the future development of Salicornia farms.

## Materials and Methods

### samples collection

The previously collected records by the Agricultural Biotechnology Research Institute of Iran (ABRII) were used to identify distribution localities for sampling [27]. Then field surveys were done to collect samples from populations throughout the natural distribution of *Salicornia* sp. Twenty-six, different geographical locations were included in this research (Fig 1). The number of plants per each location ranged from, 2 to 5 (Table 1).

**Fig 1.**
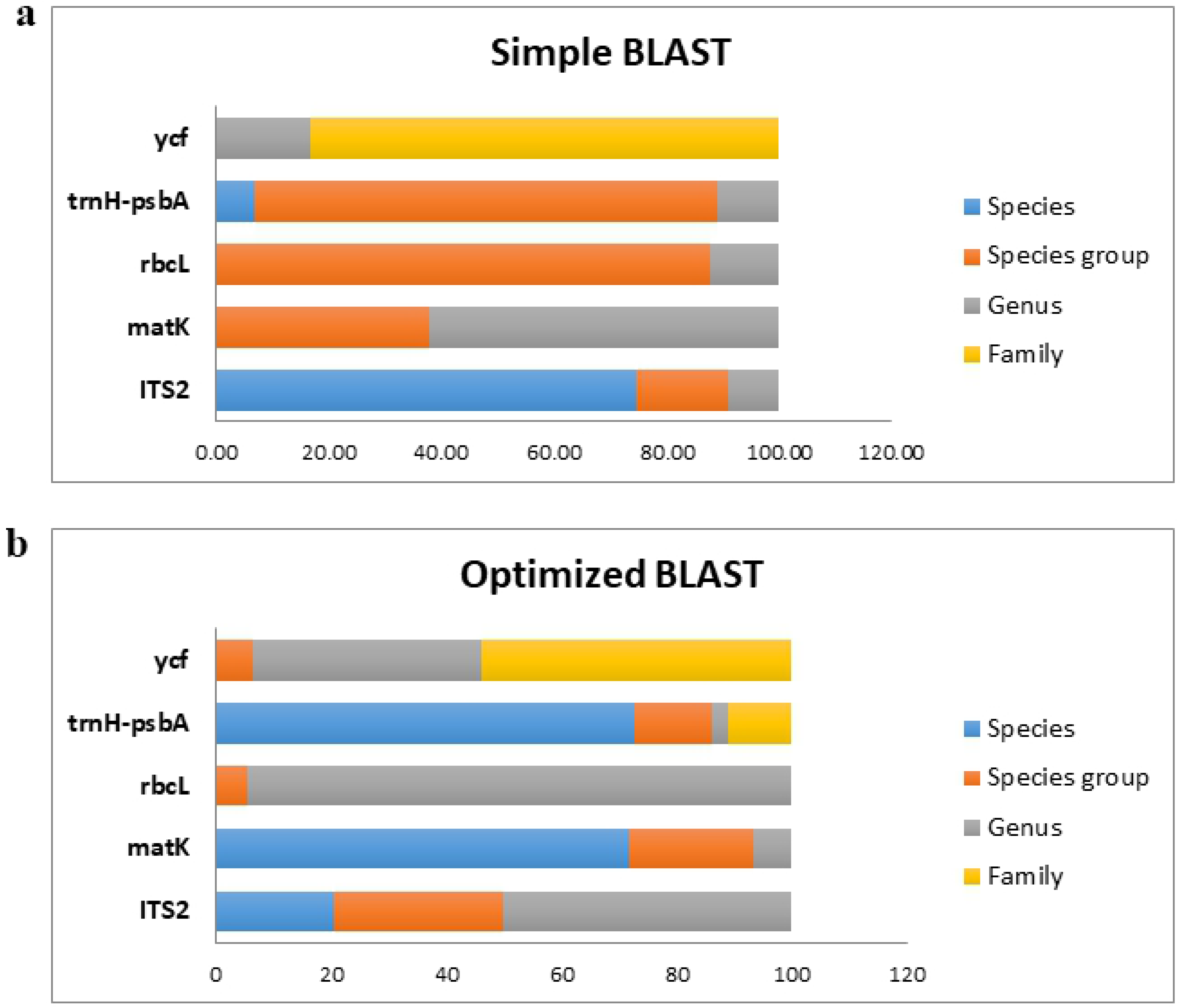
Distribution map of *Salicornia* germplasms in Iran. Maps were generated through tools in Google Earth program version 7.1.7.2606 (www.google.com/earth), based on the recorded GPS coordinates on the collection sites.

**Table 1.** Geographic information of *Salicornia* populations that were collected in this study.

| No. | Province | Collection Site | Longitude | Latitude |
| --- | --- | --- | --- | --- |
| 1 | Alborz | Jafar Abad | 50° 49' 26.511" | 35° 42' 38.401 " |
| 2 | Alborz | Eshtehard | 50° 30' 20.195" | 35° 44' 45.052 " |
| 3 | Markazi | Meyghan | 49° 48' 56.527" | 34° 10' 12.262 " |
| 4 | Markazi | Delijan | 50° 49' 58.401" | 33° 48' 38.401 " |
| 5 | Isfahan | Varzaneh | 52° 39' 53.002" | 32° 25' 22.763 " |
| 6 | Mazandaran | Chapak roud | 52° 51' 48.51" | 36° 43' 9.602 " |
| 7 | West-Azerbaijan | Gulmanxana | 45° 16' 17.644" | 37° 35' 13.912 " |
| 8 | West-Azerbaijan | Talatappe | 45° 13' 27.699" | 37° 44' 54.563 " |
| 9 | Qom | Qom highway | 51° 10' 43.625" | 35° 19' 18.318 " |
| 10 | Sistan & Blaochestan | Chabahar | 60° 40' 20.423" | 25° 17' 21.001 " |
| 11 | Sistan & Blaochestan | Gwadr | 62° 22' 29.560" | 25° 20' 6.936 " |
| 12 | Tehran | Robat Karim | 50° 57' 44.412" | 35° 26' 37.730 " |
| 13 | Tehran | Varamin | 51° 17' 40.124" | 34° 56' 53.952 " |
| 14 | Golestan | Bandar Torkamen | 54° 3' 0.072" | 36° 53' 18.160 " |
| 15 | Golestan | Aq Qala | 54° 22' 46.480" | 37° 5' 58.844 " |
| 16 | Boshehr | Dayyer | 51° 58' 31.501" | 27° 50' 31.642 " |
| 17 | Boshehr | Helleh | 50° 49' 52.812" | 29° 11' 22.106 " |
| 18 | East-Azerbaijan | Nazarkahrizi | 46° 54' 52.339" | 37° 21' 34.416 " |
| 19 | East-Azerbaijan | Qushchi | 45° 4' 39.946" | 37° 59' 40.941 " |
| 20 | East-Azerbaijan | Sharafkhaneh | 45° 28' 33.041" | 38° 10' 2.236 " |
| 21 | East-Azerbaijan | SarayDeh | 45° 38' 48.551" | 37° 52' 47.262 " |
| 22 | East-Azerbaijan | Qareh Qeshlaq | 45° 58' 6.110" | 37° 13' 46.699 " |
| 23 | Fars | Tashk | 53° 35' 41.686" | 29° 49' 15.232 " |
| 24 | Fars | Maharlu | 53° 45' 50.043" | 29° 31' 45.922 " |
| 25 | Kerman | Nogh | 56° 0' 0219" | 30° 39' 17.028 " |
| 26 | Kerman | Shahrbabak | 54° 39' 56.807" | 30° 6' 8.463 " |

### Molecular analysis

#### DNA extraction and barcoding

DNA extraction was carried out using DNA extraction kit (Core Bio, South Korea). The quality and quantity of DNA was determined using 0.8% agarose gel and NanoDrop ND-1000 spectrophotometer (Nanodrop Technologies, Wilmington, DE, USA), respectively. DNA barcoding was carried out using core-barcode consisting of two plastid coding regions, (*rbcL*+*matK*) and supplemented with one non-coding intergenic spacer (*trnH-psbA*) from the chloroplast genome, one internal transcribed spacer (ITS2) from the nuclear genome and one plastid coding region (*ycf*) [26,28] (see S1 Table). For all markers, 50 μl polymerase chain reactions (PCRs) were carried out in a Veriti ABI thermal cycler (ABI Inc, USA) using 1 U Taq polymerase (Invitrogen or Promega), 2.5 μl 10 Taq buffer, 1.5 μl MgCl_2_ (25 mM), 0.5 μl dNTPs (10 mM), 0.35 μl primers (20 pmol), 1.0 μl template DNA and 18.85 μl dH_2_O. Then, the PCR products were purified and directly sequenced in both directions to minimize PCR artifacts, ambiguities, and base-calling errors on an automated ABI Prism 3730 XL Genetic Analyzer machine using ABI BigDye v3.1 Terminator Sequencing chemistry (Macrogen Inc, South Korea). However, the samples with high GC content were cloned and then sequenced. The sequencing results were then sorted and trimmed using FinchTV software and analyzed by BLASTP, BLASTN, and LALIGN software. Finally, DNA barcode data compared to GenBank or otherwise publicly available in the BOLD database based on different integration strategies (coding, non-coding, plastid, and total combination) using a similarity-based method. Briefly, the sequence similarity-based method was utilized for species identification, using simple and optimized BLAST, for different marker integration strategies (single, coding, non-coding, plastid, and total combination). First, sequences were queried using megablast at Geneious software and NCBI BLASTn against the nucleotide database. Then, a simple method was considered the top 10 hit, and the top 100 records were used to put extra weight on the identity value using “max score*(query cover/identity)” formula on an optimized method. Finally, taxonomic identifications were allocated based on the combination of the identity score (High identity: X≥95%; Medium identity: 90% ≤ X ≤ 95%; Low identity: X ≤ 90%) and the number of species within 1% deviation of the calculated similarity score (S1 Fig). Plant Sequences database (BOLD systems) and ITS2 database (V5) were also used as a validation tool for the core barcodes (matK, rbcL) and ITS2 [29,30].

### Analysis of genetic diversity

Analysis of molecular variance (AMOVA) was carried out using Arlequin version 3.5 [31]. The significance level of *F*_*st*_ statistics was also performed using the software by a nonparametric permutation procedure with 1023 randomizations. Statistical calculations and graphics for *F*_*st*_ were conducted using Arlequin and R version 3.6.1 (cran.r-project.org) [32]. Further, the online version of Automatic Barcode Gap Discovery (ABGD) was used to generate the genetic distance histograms and ranked distance based on K2P distance (https://bioinfo.mnhn.fr/abi/public/abgd/abgdweb.html[33].

### Species distribution modeling

The modeling program MAXENT was used to model Iranian *Salicornia* sp. potential distribution based on an algorithm of maximum entropy to calculate the ecological niche of species and to find out the potential natural distribution of areas and to limit climatic factors [34]. The web-based platform, WorldClim database (http://www.diva-gis.org/Data) version 1.3, October 2004 [35], which includes major climate databases from different sources, was used to provide nineteen environmental variables (BIOCLIM). Then, environmental data were added from the FAO Map on Global Ecological Zones [36], to build up the potential natural distribution model. The Maximum threshold sensitivity plus specificity was = 0.180 for predicting species geographic distribution (for details see Liu et al. [37]).

Further, the geographic observations used for modeling of the potential distribution of *Salicornia sp*. were first filtered, and outliers were then detected in DIVA-GIS (www.diva-gis.org) [38]. Then, all occurrence records and extreme values (minimum of 3 out of 19 bioclimatic variables examined) were rechecked for the inconsistency of their coordinates with administrative area level 1 and their climatic parameters based on the Reverse Jackknife method [39]. Finally, the constructed model assessed with the area under the receiver operating characteristic (ROC) curve [40]; the area under the curve (AUC) statistics as an independent threshold, measures the model performance, ranging from 0.5 to 1.0 [41].

## Results and Discussion

The success rates of PCR amplification and sequencing of the five barcode markers along with their phylogenetic tree using signle barcodes are shown in Fig 2 and S1 File. The three genomic and plastid DNA regions, including ITS2, *trnH*-*psbA* spacers and *ycf* genes and a combination of *matK*-*rbcL* were used as acceptable standard barcodes. Among the markers tested, *rbcL* had the highest amplification and recovery rates (98.90%), followed by *trnH*-*psbA* (82.42%), *matK* (80.21%), *ycf* (69.23%) and the rate for ITS2 was the lowest (65.93%). A low percentage amplification rate and recovery for ITS2 were due to the lack of suitable public primer, the incongruence of multiple copies, and or technical problems. In total, the sequence alignment method comprised of 60 ITS2 sequences, 73 *matK* sequences, 90 sequences of *rbcL*, 75 sequences of *trnH-psbA,* and 63 sequences of *ycf*.

**Fig 2.**
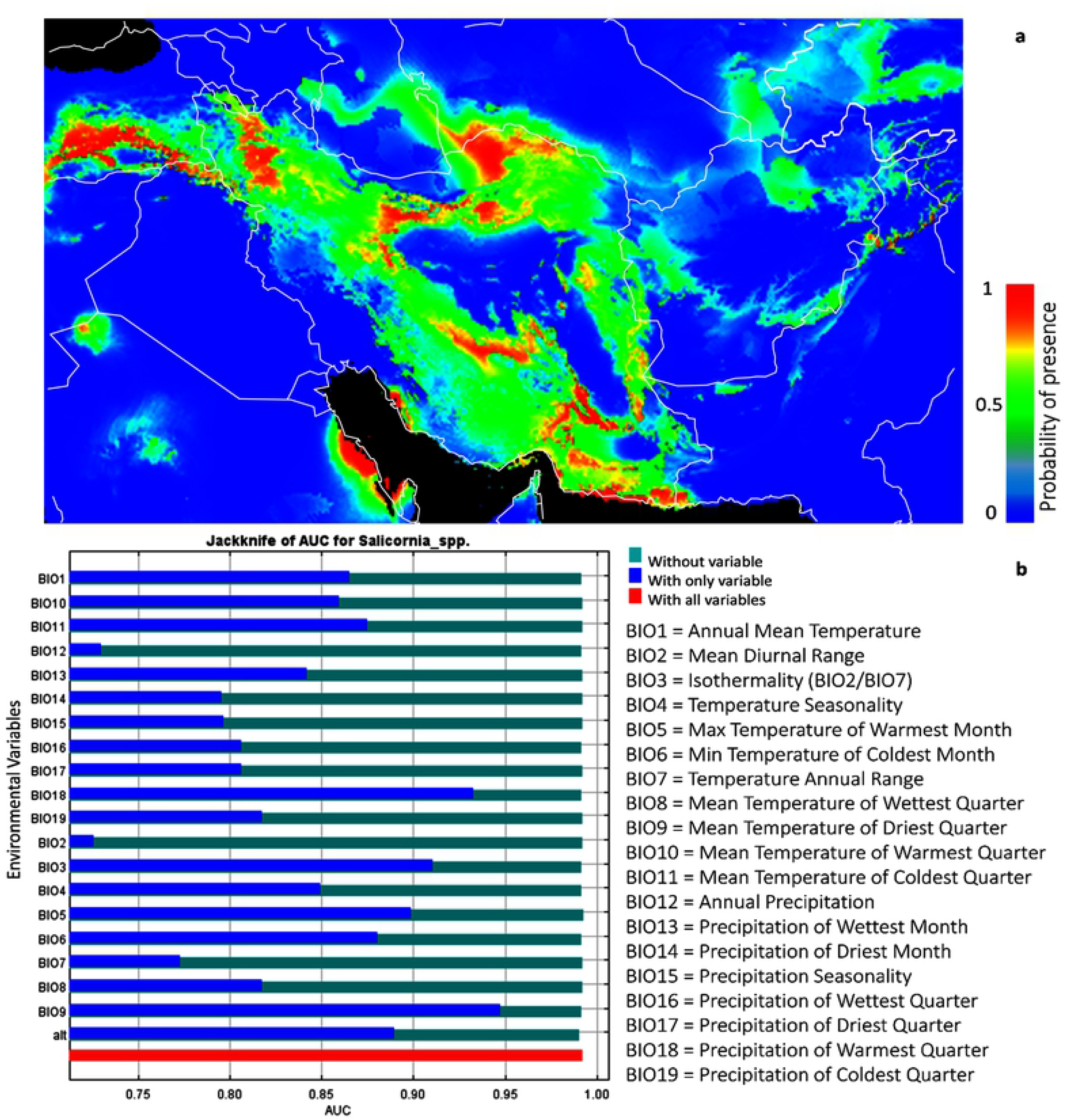
The success rates of PCR amplification and sequencing of the five-barcode fragment and 26 different geographical locations.

Simple and optimized BLAST was carried for each of these sequences based on NCBI and ITS database. The efficiency of every single marker, as well as their combination for taxonomic identification, is presented in Fig 3. The highest level of identification using the single marker and simple BLAST at the species level, species group, genus, and family groups were obtained for ITS2 (74.55%), *rbcL* (98.78%), *matK* (62.19%) and *ycf* (83.08%) respectively while the discrimination efficiency using optimized BLAST was 70% (*matK* and *trnH*-*psbA*), 29.63% (ITS2), 94.44% (*rbcL*) and 53.96% (*ycf*) respectively (Fig 3). Likewise, data integration showed a higher identification at the level of the species group [42]. This group included *S. persica, S. europea, S. patula, S. brachiate, S. herbacea* and *S. maritime* suggesting species such as *S. europea, S. patula* and *S. herbacea* are similar or synonymous (The Plant List (2013). Version 1.1. Published on the Internet; http://www.theplantlist.org/ (accessed 1st January)). Automatic barcode gap discovery (ABGD) analysis detected a barcoding gap between the intraspecific and interspecific distance of barcode markers. Two distinct groups were classified by ABGD analysis for all barcode markers. In the ABGD system, specimens identified as one group could be considered as one species [33].

**Fig 3.**
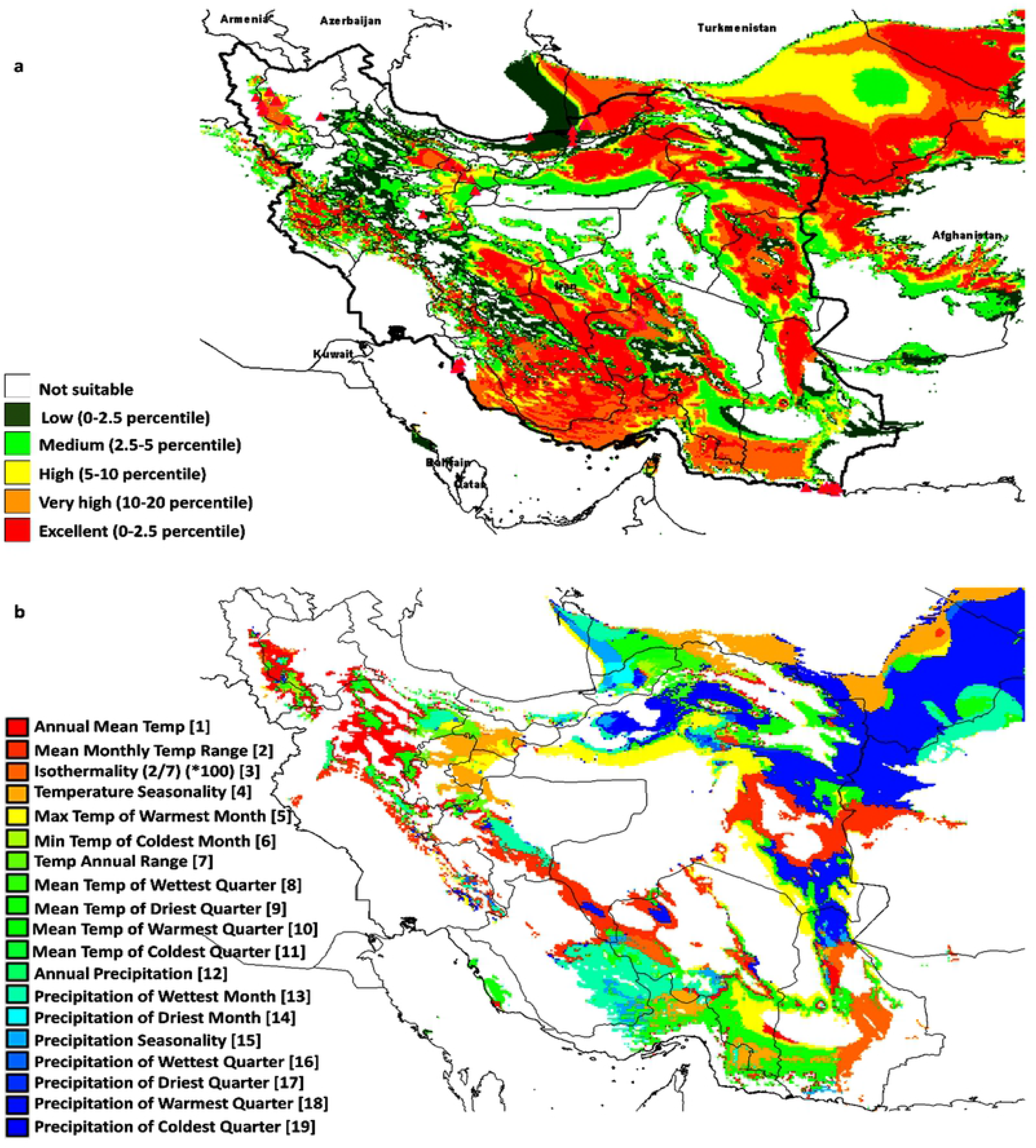
Comparison of the efficiency of DNA barcoding using selected regions; a) simple and b) optimized BLAST based on NCBI and ITS database.

It should be noted that the ABGD algorithm yielded the same results when applied in the three implemented models (JC, K2P, and Simple Distance). Histograms of sequence divergence values and ranked distances among barcode sequences in Salicornia complex are shown in S2 Fig. A combination of barcode markers in noncoding, coding, core, and cpDNA group did not increase discrimination success. However, it confirmed that the ABGD results suggest the existence of two Salicornia species. The highest discrimination success in all combinations obtained for the *S. europea* species followed by *S. brachiata* (Fig 4). Further, the low success rate of markers might be related to the spreading and distribution of Salicornia germplasm species in Iran. This index is inverted with the amount of intra and inter-species gene flow. According to this theory, populations with a low distribution index are geographically or physiologically related to each other. The low distribution index may cause the variants of neutral mutations to spread slowly throughout the entire population. Thus the required time for lineage sorting for each gene locus increase by the natural genetic flow. In this case, the identification of specific barcodes is difficult for each species [43].

**Fig 4.**
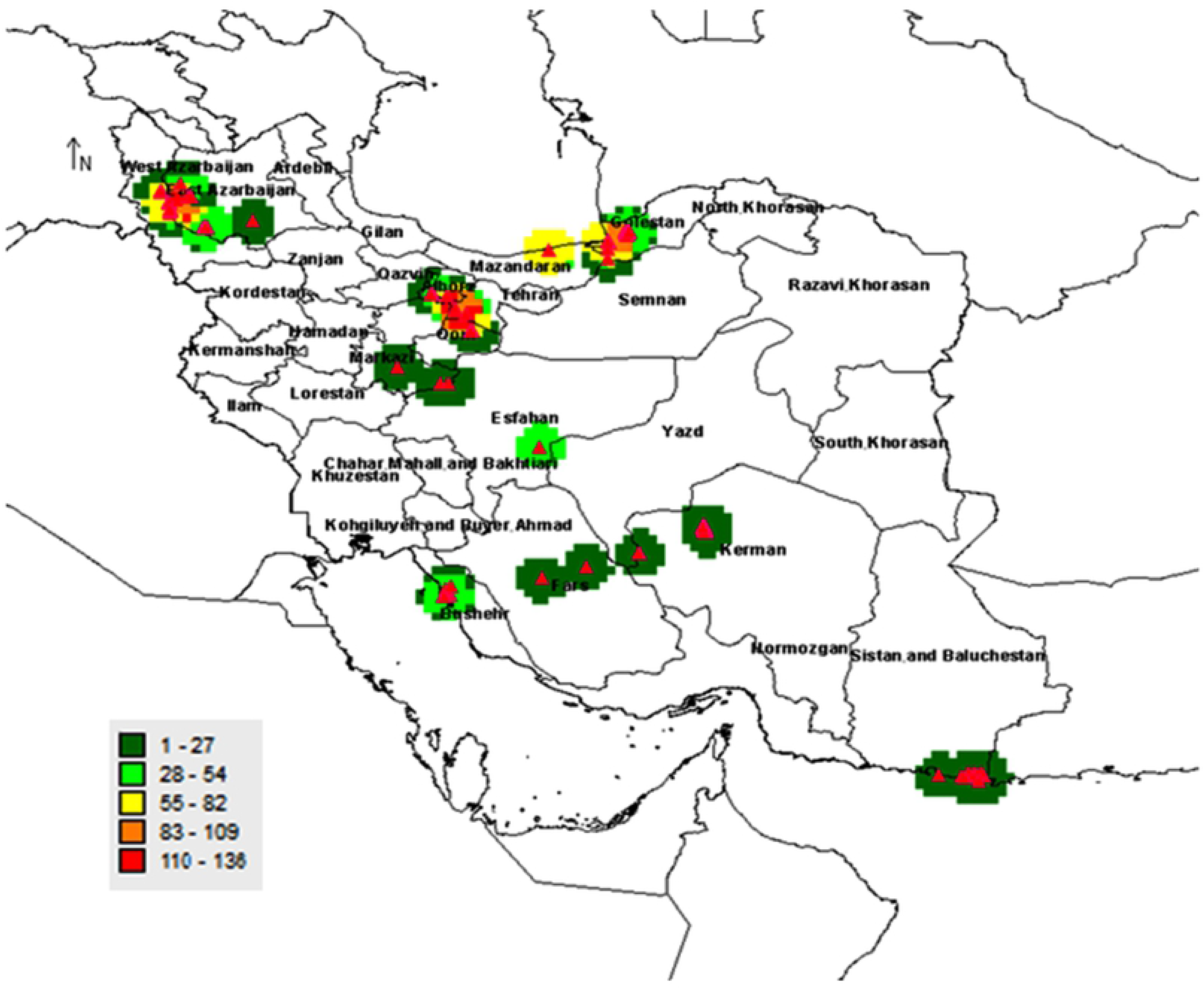
Performances of *matK*, *rbcL*, *trnH*-*psb* A, *ITS2* and *ycf* in resolving Salicornia species as different noncoding, coding, core and cpDNA combinations.

Another consequence is that an increase in the penetration coefficient of a species in the adjacent species is caused by genetic flow, affecting the efficiency rate of the DNA barcoding method. In this study, the barriers and geographic distances, self-pollination, and cleistogamy reduced the gene flow rate among the plausible species of Salicornia populations. Later, the Iranian Salicornia germplasm has been studied with respect to the different nuclear and plastid sites with different heritability patterns in each group. As previously reported, the inheritance of plastid DNA is maternal in most angiosperms. Thus plastid haplotypes disperse through seeds [44]. In contrast, nuclear variants are distributed by seed and pollen and exhibited by biparental inheritance. The expected distribution range of pollen is higher relative to seed. Thus the genetic differentiation of the populations should be higher through plastid markers [45–47]. Here, molecular variance analysis (AMOVA) and *F*_*st*_ index for core-barcode plastid (matK, rbcL) and nuclear marker (ITS2) showed that molecular variance within populations of the same species is greater than between populations indicating a higher genetic diversity (Fig 5). The largest difference between and within populations was observed in Dayer and Shahrbabbak genotypes based on ITS2 and matK markers, respectively. However, the rbcL fragment was not suitable for differentiation between populations. The highest Nei’s distance was obtained in Golmanxana and Qaregheshlaq genotypes based on ITS2 and rbcL markers. Moreover, the highest Fst index was observed in the Golmanxana genotype for the ITS2 marker, suggesting a low genetic flow rate compared to other genotypes. A higher Fst index conserves the allelic diversity needed for the conservation and exploitation of genetic resources. Further, the distribution of seed and pollen varies among populations, and in most cases, the genetic differentiation between populations depends on the geographical barriers and distance [48].

**Fig 5.**
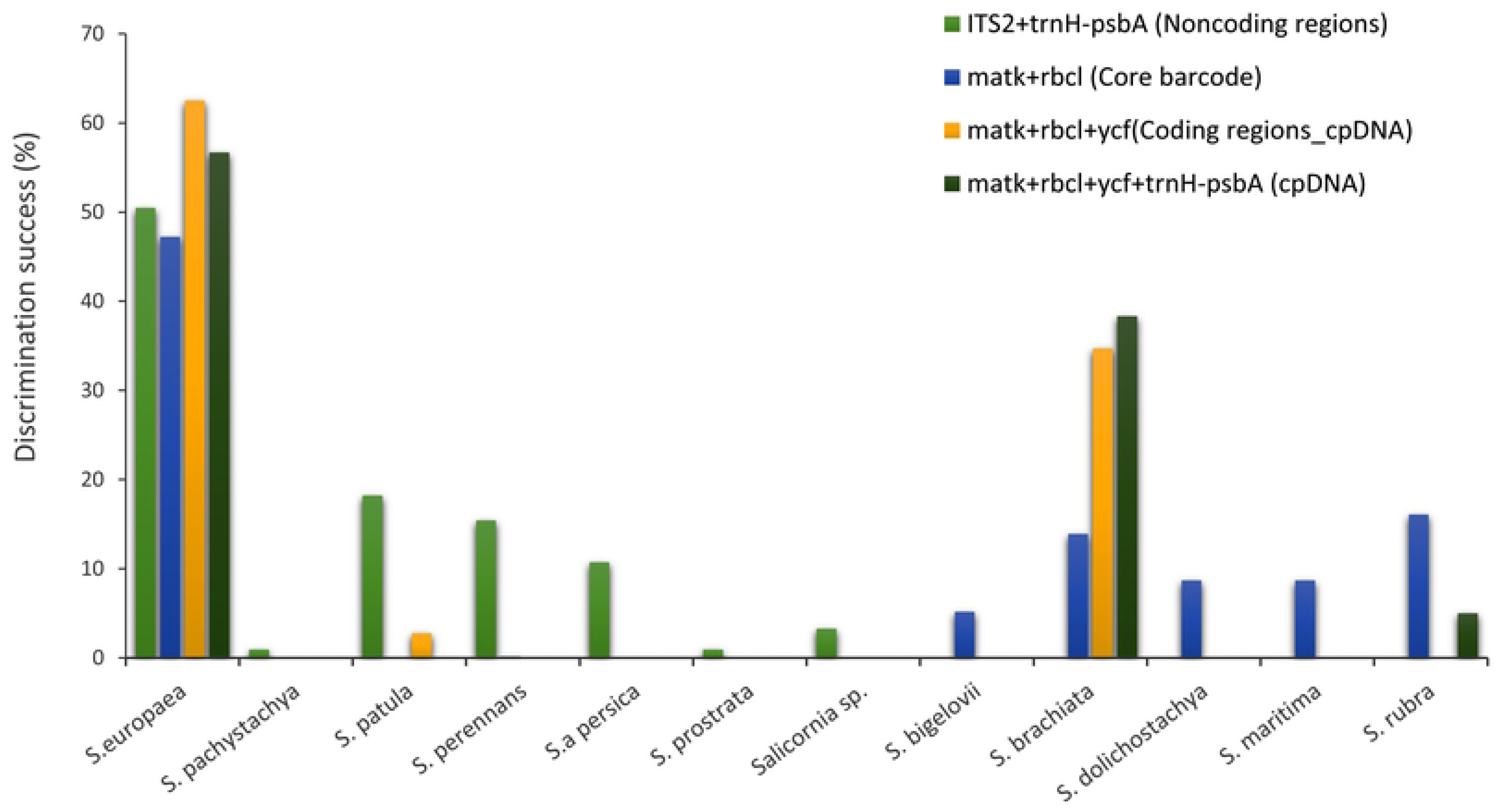
Inter and intra molecular variance analysis and*F*_*st*_ index in*Salicornia* germplasm in Iran, based on plastid and nuclear markers.

The spatial analysis helps in better understanding and more accurate identification of biodiversity and developing strategies for identifying, managing and exploiting genetic resources [49]. Outputs then provide vital information for prioritizing of protection and the Agro-Ecological Areas [50] finding. This finding may suggest that the new species can be grown in an environment outside its endemic region if additional resources such as water or soil nutrients are available. It should be noted that combining spatial results with other data is very useful in the management and exploitation of genetic reserves more efficiently [51]. The basis of spatial analysis in biodiversity is observed data from the sampled collection areas. In our study the geographical data included identification codes, taxonomic names, geographic characteristics, and sampling locations (Fig 6). This information is commonly used for spatial diversity and distribution analysis. Biodiversity of plants is studied at three levels comprising of species-level and genetic level in a population (ecosystem). Here, Salicornia was studied at the species level or alpha diversity. In this case, the species level was observed in species diversity and evaluated in being absent or present conditions in each specific area.

**Fig 6.**
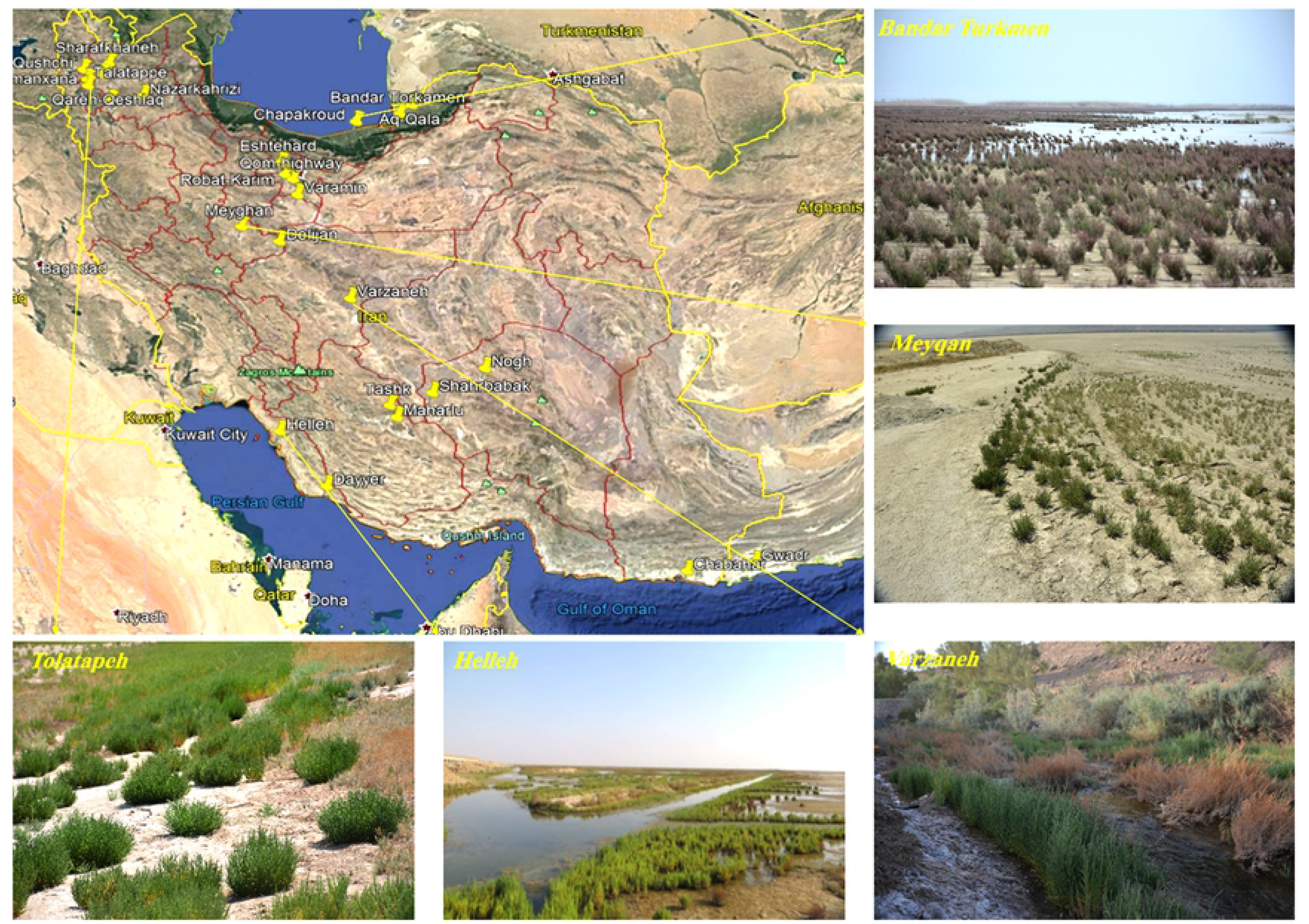
Distribution of the*Salicornia* populations in Iran based on the sampling areas. The regions with the highest and the lowest number of samples are depicted in red and dark green, respectively. The map was created using Google Earth program version 7.1.7.2606 (www.google.com/earth).

Further, the determination of diversity (including species, genotype, or ecotype) in different subunits was one of the most crucial aims in this study. The sub-units were areas that Salicornia species were found in a previous study. Consequently, diversity was studied at the alpha level, and the sampling areas were mapped using the observed number for each sample and their distributions (Fig 7). The regions were then classified into five groups based on their distributions. The area with the highest and the lowest number of samples is depicted in red and dark green, respectively, in Fig 7. For example, if a plant protection program is considered, the areas with the highest alpha diversity (red) are in the top priority. In the current study, primary data in specific formats were prepared for DIVA and MAXENT software; then, the climate data information was converted into a particular format for the software, and 19 different climate variables affecting the distribution of species were evaluated [52] (Fig 8 and S2 File). As shown in Fig 8, isothermality (diurnal range mean/temperature annual range) *100, mean temperature of driest quarter, precipitation of warmest quarter, the maximum temperature of the warmest month, altitude, temperature seasonality, mean temperature of coldest quarter, minimum temperature of the coldest month, and annual mean temperature were the most critical variables to the Salicornia MAXENT model, based on jackknife test AUC. The average test AUC for each replicate runs were 0.994. Likewise, the standard deviation was 0.002.

**Fig 7.**
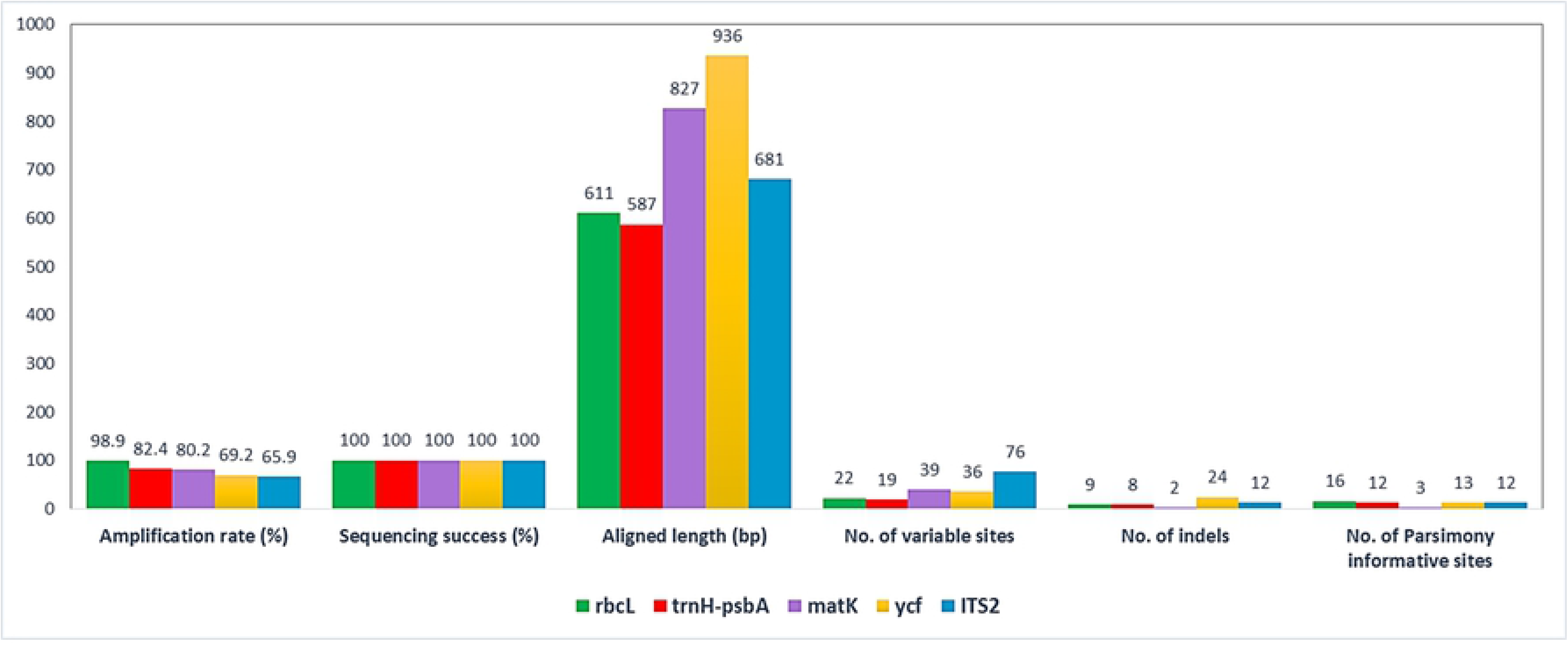
Geographical distribution of*Salicornia* in Iran generated by MAXENT software (a); the probability of presence increase from blue to red, (b) the effect of climates parameters on*Salicornia* distribution in Iran. The map was created using free open source MAXENT software version 3.4.1 (https://biodiversityinformatics.amnh.org/open_source/maxent/).

**Fig 8.**
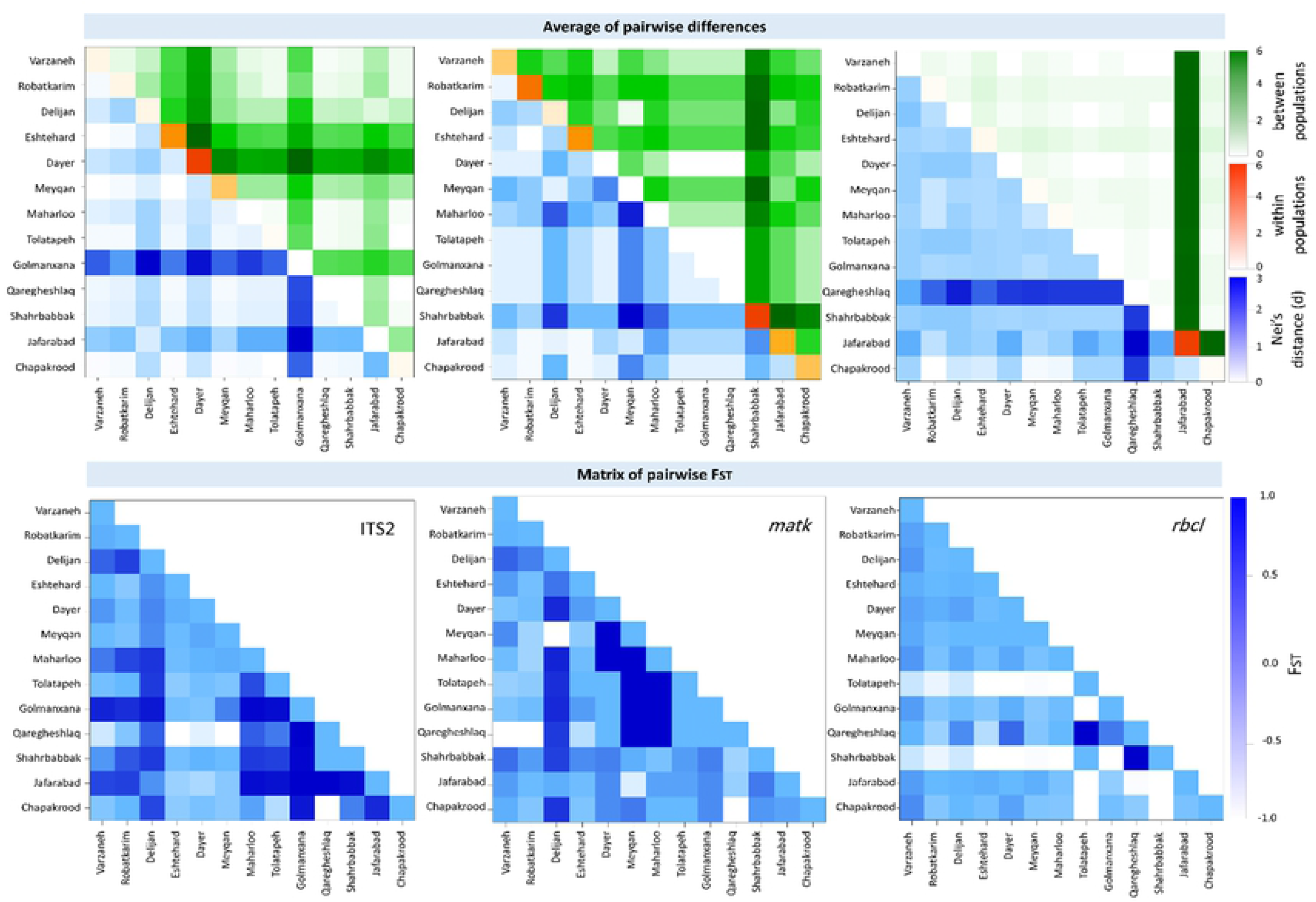
Prediction model of*Salicornia* distribution (a) and limiting factors (b), based on geographical data and cluster analysis of sampled material. Different niches for protection and cultivation of Salicornia was shown with red to green color. The most competence areas for natural identification or cultivation of Salicornia when optimum water and soil parameter conditions are available shown in red colors while dark green colors show the least potential areas. The map was created using free software DIVA-GIS Version 7.5 (www.diva-gis.org).

We also studied ecological niches on the conservation and utilization of genetic resources. This concept has been used to determine the priority of areas suitable for the conservation of wild species or the banking of natural genetic resources. Ecological niches are occupied area by a species in natural environmental conditions. Initial niches were also identified in this study. These areas have favorable climatic conditions for Salicornia growth and can be used to found these species.

Further, in the field trial condition, the coverage of the primary and ecological niches was confirmed. However, in some primary niches, Salicornia samples were not found that may be due to the negative effect of climates or ecological parameters. Further, Geographic Information Systems (GIS) was used to model ecosystem nets based on available environmental data of each collected samples [6]. This model was drawn using DIVA and MAXENT software (Figure 8). The map showed different niches for the protection and cultivation of Salicornia. The most competence areas for natural identification or cultivation of Salicornia shown are red colors when optimum water and soil parameters condition are available. However, the dark green colors show the least potential regions. Together, the model exhibited saline soil, and inefficient land might be used for industrial cultivation of Salicornia in Iran. Murray‐smith et al. (2009), previously applied MAXENT modeling and DIVA-GIS analysis to identify priority areas for the conservation of Myrtaceae. Their model showed observed species occurrences and predicted species occurrences and indicated of complementarity analysis congruent in identifying areas with the most endemic species.

## Conclusion

We validated genes in the field of DNA barcoding in Salicornia plants using matK, rbcL, trnH-psbA, ycf and ITS2, identifying species groups and the possible model of gene flow within and among Salicornia population. Among the genetic markers tested, *rbcL* had the highest amplification and recoverability rates (98.90%), followed by *trnH*-*psbA* (82.42%), *matK* (80.21%), *ycf* (69.23%) and the rate for ITS2 was the lowest (65.93%). Data integration showed identification at the level of the species group was higher. This group included *S. persica, S. europea, S. patula, S. brachiate, S. herbacea,* and *S. maritime*. Molecular variance analysis and *F*_*st*_ index for plastid and nuclear markers showed genetic differences within the population of the same species is greater than between the populations indicating a high genetic diversity among the genotypes of each population. We also studied ecological niches on conservation and utilization of genetic resources, to determine the priority of areas suitable for conservation of wild species or banking of natural genetic resources. Our results provided valuable information on the diversity of specific geographical regions, conservation status of existing species, prioritization of conservation areas, and selection of regions for Agro-Ecological, which might be led to the development of industrial agriculture.

## Supporting information

**Fig S1. Taxonomic identifications based on the combination of the identity score value. (High identity: X≥95%; Medium identity: 90% ≤ X ≤ 95%; Low identity: X ≤ 90%).**

**Fig S2. Automatic Barcode Gap Discovery (ABGD) analysis of barcoding markers, (a) Histogram of distances, (b) graph of ranked distances, (c) Automatic partition.**

**Table S1. DNA barcode primers sequences used in this study.**

**S1 File. The phylogenetic tree using a single fragment of ITS2, matK, rbcl, trn and ycf.**

**S2 File. The primary data used in DIVA and MAXENT software.**

## References

1. Bartels D, Sunkar R. Drought and Salt Tolerance in Plants. CRC Crit Rev Plant Sci. 2005;24: 23–58. doi:10.1080/07352680590910410

2. Godfray HCJ, Beddington JR, Crute IR, Haddad L, Lawrence D, Muir JF, et al. Food security: the challenge of feeding 9 billion people. Science (80-). 2010;327: 812–8. doi:10.1126/science.1185383

3. Flowers TJ, Hajibagheri MA, Clipson NJW. Halophytes. Q Rev Biol. 1986;61: 313–337. doi:10.1086/415032

4. Flowers TJ. Improving crop salt tolerance. J Exp Bot. 2004;55: 307–319. doi:10.1093/jxb/erh003

5. Hadi MR, Khiyam-Nekoie SM, Khavarinejad R, Khoshkholgh Sima NA, Yavari P. Accumulation and role of ions (Ca2+, Mg2+, SO4-2) on salt tolerance in Triticum turgidum L. . J Biol Sci. 2008;8: 143–148. Available: https://scholar.google.com/scholar?cluster=16416560739638406201&hl=en&oi=scholarr

6. Kadereit G, Ball P, Beer S, Mucina L, Sokoloff D, Teege P, et al. A taxonomic nightmare comes true: phylogeny and biogeography of glassworts (Salicornia L., Chenopodiaceae). Taxon. 2007;56: 1143–1170. Available: https://www.ingentaconnect.com/contentone/iapt/tax/2007/00000056/00000004/art00013?crawler=true

7. Alexandratos N, Bruinsma J. World agriculture towards 2030/2050: the 2012 revision. ESA Working Paper No. 12-03. Rome: Food and Agriculture Organization, Agricultural Development, Economics Division; 2012. Available: https://ageconsearch.umn.edu/record/288998/

8. Teege P, Kadereit JW, Kadereit G. Tetraploid European Salicornia species are best interpreted as ecotypes of multiple origin. Flora - Morphol Distrib Funct Ecol Plants. 2011;206: 910–920. doi:10.1016/J.FLORA.2011.05.009

9. Flowers TJ, Colmer TD. Salinity Tolerance in Halophytes. New Phytol. 2008;179: 945–963. doi:10.2307/25150520

10. Khosh Kholgh Sima NA, Tale Ahmad S, Pessarakli M. Comparative study of different salts (sodium chloride, sodium sulfate, potassium chloride, and potassium sulfate) on growth of forage species. J Plant Nutr. 2013;36: 214–230. doi:10.1080/01904167.2012.739242

11. Öztürk M, Güvensen A, Sakçali S, Görk G. Halophyte plant diversity in the Irano-Turanian phytogeographical region of Turkey. Biosaline Agriculture and High Salinity Tolerance. Basel: Birkhäuser Basel; 2008. pp. 141–155. doi:10.1007/978-3-7643-8554-5_14

12. Flowers TJ, Galal HK, Bromham L. Evolution of halophytes: multiple origins of salt tolerance in land plants. Funct Plant Biol. 2010;37: 604. doi:10.1071/FP09269

13. Hammer K, Khoshbakht K. Towards a ‘red list’ for crop plant species. Genet Resour Crop Evol. 2005;52: 249–265. doi:10.1007/s10722-004-7550-6

14. Eganathan P, SR Subramanian HM, Latha R, Rao CS. Oil analysis in seeds of Salicornia brachiata. Ind Crops Prod. 2006;23: 177–179. doi:10.1016/J.INDCROP.2005.05.007

15. Ventura Y, Wuddineh WA, Shpigel M, Samocha TM, Klim BC, Cohen S, et al. Effects of day length on flowering and yield production of Salicornia and Sarcocornia species. Sci Hortic (Amsterdam). 2011;130: 510–516. doi:10.1016/J.SCIENTA.2011.08.008

16. Katschnig D, Broekman R, Experimental JR-E and, 2013 undefined. Salt tolerance in the halophyte Salicornia dolichostachya Moss: Growth, morphology and physiology. Environ Exp Bot. 2013;92: 32–42. Available: http://www.sciencedirect.com/science/article/pii/S0098847212000895

17. Glenn EP, Anday T, Chaturvedi R, Martinez-Garcia R, Pearlstein S, Soliz D, et al. Three halophytes for saline-water agriculture: An oilseed, a forage and a grain crop. Environ Exp Bot. 2013;92: 110–121. doi:10.1016/J.ENVEXPBOT.2012.05.002

18. Liu X, Xia Y, Wang F, Sun M, Jin Z, Science GW-F, et al. Analysis of Fatty Acid Compositions of Salicornia Europaea L. Seed Oil. Food Sci. 2005;2. Available: http://en.cnki.com.cn/Article_en/CJFDTOTAL-SPKX200502042.htm

19. Muscolo A, Panuccio MR, Piernik A. Ecology, Distribution and Ecophysiology of Salicornia Europaea L. Springer, Dordrecht; 2014. pp. 233–240. doi:10.1007/978-94-007-7411-7_16

20. Petit RJ, Duminil J, Fineschi S, Hampe A, Salvini D, Vendramin GG. INVITED REVIEW: Comparative organization of chloroplast, mitochondrial and nuclear diversity in plant populations. Molecular Ecology. John Wiley & Sons, Ltd (10.1111); 2004. pp. 689–701. doi:10.1111/j.1365-294X.2004.02410.x

21. Kadereit G, Piirainen M, Lambinon J, Vanderpoorten A. Cryptic taxa should have names: Reflections in the glasswort genus Salicornia (Amaranthaceae). Taxon. 2012;61: 1227–1239. Available: https://www.ingentaconnect.com/contentone/iapt/tax/2012/00000061/00000006/art00005?crawler=true

22. Kaur G, Kumar S, Nayyar H, Upadhyaya HD. Cold Stress Injury during the Pod-Filling Phase in Chickpea (*Cicer arietinum* L.): Effects on Quantitative and Qualitative Components of Seeds. J Agron Crop Sci. 2008;194: 457–464. doi:10.1111/j.1439-037X.2008.00336.x

23. Vorasoot N, Songsri P, Akkasaeng C, Jogloy S, Patanothai A. Effect of water stress on yield and agronomic characters of peanut (Arachis hypogaea L.). J Sci Technol. 2003;25: 283–288. Available: https://www.researchgate.net/profile/Chutipong_Akkasaeng/publication/26483170_Effect_of_water_stress_on_yield_and_agronomic_characters_of_peanut_Arachis_hypogaea_L/links/0c96051526ae30ecd8000000/Effect-of-water-stress-on-yield-and-agronomic-characters-of-

24. Ingrouille MJ, Pearson J. The pattern of morphological variation in the Salicornia europaea L. aggregate (Chenopodiaceae). Watsonia. 1987;16: 269–281. Available: http://archive.bsbi.org.uk/Wats16p269.pdf

25. Neves SS, Forrest LL. Plant DNA Sequencing for Phylogenetic Analyses: From Plants to Sequences. Humana Press; 2011. pp. 183–235. doi:10.1007/978-1-61779-276-2_10

26. Hollingsworth PM, Graham SW, Little DP. Choosing and Using a Plant DNA Barcode. Steinke D, editor. PLoS One. 2011;6: e19254. doi:10.1371/journal.pone.0019254

27. Reiahisamani N, Esmaeili M, Khoshkholgh Sima NA, Zaefarian F, Zeinalabedini M. Assessment of the oil content of the seed produced by Salicornia L., along with its ability to produce forage in saline soils. Genet Resour Crop Evol. 2018;65: 1879–1891. doi:10.1007/s10722-018-0661-2

28. Dong W, Liu J, Yu J, Wang L, Zhou S. Highly Variable Chloroplast Markers for Evaluating Plant Phylogeny at Low Taxonomic Levels and for DNA Barcoding. Moustafa A, editor. PLoS One. 2012;7: e35071. doi:10.1371/journal.pone.0035071

29. Kearse M, Moir R, Wilson A, Stones-Havas S, Cheung M, Sturrock S, et al. Geneious Basic: An integrated and extendable desktop software platform for the organization and analysis of sequence data. Bioinformatics. 2012;28: 1647–1649. doi:10.1093/bioinformatics/bts199

30. Ghorbani A, Saeedi Y, de Boer HJ. Unidentifiable by morphology: DNA barcoding of plant material in local markets in Iran. Bussmann R, editor. PLoS One. 2017;12: e0175722. doi:10.1371/journal.pone.0175722

31. Excoffier L, Lischer HE. Arlequin suite ver 3.5: a new series of programs to perform population genetics analyses under Linux and Windows. Mol Ecol Resour. 2010;10: 564–567. doi:10.1111/j.1755-0998.2010.02847.x

32. Weir BS, Cockerham CC. Estimating *F* - statistics for the analysis of population structure. Evolution (N Y). 1984;38: 1358–1370. doi:10.1111/j.1558-5646.1984.tb05657.x

33. Puillandre N, Lambert A, Brouillet S, Achaz G. ABGD, Automatic Barcode Gap Discovery for primary species delimitation. Mol Ecol. 2012. doi:10.1111/j.1365-294X.2011.05239.x

34. Warren DL, Seifert SN. Ecological niche modeling in Maxent: the importance of model complexity and the performance of model selection criteria. Ecol Appl. 2011;21: 335–342. doi:10.1890/10-1171.1

35. Escudero A, Iriondo JM, Torres ME. Spatial analysis of genetic diversity as a tool for plant conservation. Biol Conserv. 2003;113: 351–365. doi:10.1016/S0006-3207(03)00122-8

36. MacDicken K, Jonsson Ö, Piña L, Maulo S, Contessa V. Global forest resources assessment 2015: how are the world’s forests changing? 2016 [cited 30 Jul 2019]. Available: http://agris.fao.org/agris-search/search.do?recordID=XF2017001127

37. Liu C, Berry PM, Dawson TP, Pearson RG. Selecting thresholds of occurrence in the prediction of species distributions. Ecography (Cop). 2005;28: 385–393. doi:10.1111/j.0906-7590.2005.03957.x

38. Hijmans RJ, Guarino L, Cruz M, Rojas E. Computer tools for spatial analysis of plant genetic resources data: 1. DIVA-GIS. Plant Genetic Resources Newsletter. 2001. Available: https://books.google.com/books?hl=en&lr=&id=6QdBUMOtx0YC&oi=fnd&pg=PA15&dq=Computer+tools+for+spatial+analysis+of+plant+genetic+resources+data:+1.+DIVA-GIS.+Plant+Genetic+Resources+Newsletter&ots=pTiot0bO9v&sig=uK8ZvGqhL84JF125O2r0-meCbNE

39. Chapman A. Principles and methods of data cleaning. GBIF; 2005. Available: https://books.google.com/books?hl=en&lr=&id=44gDJTEJoVIC&oi=fnd&pg=PA1&dq=Reverse+Jackknife+method+(Chapman+2005&ots=vSlT4ce3mp&sig=o8vwrA8B135r3bRtzzNwspnn05s

40. Nyfløt LT, Sandven I, Stray-Pedersen B, Pettersen S, Al-Zirqi I, Rosenberg M, et al. Risk factors for severe postpartum hemorrhage: A case-control study. BMC Pregnancy Childbirth. 2017;17: 1–9. doi:10.1186/s12884-016-1217-0

41. Phillips SJ, Anderson RP, Schapire RE. Maximum entropy modeling of species geographic distributions. Ecol Modell. 2006;190: 231–259. doi:10.1016/J.ECOLMODEL.2005.03.026

42. Moritz C, Cicero C. DNA Barcoding: Promise and Pitfalls. Charles Godfray, editor. PLoS Biol. 2004;2: e354. doi:10.1371/journal.pbio.0020354

43. Elith J, Phillips SJ, Hastie T, Dudík M, Chee YE, Yates CJ. A statistical explanation of MaxEnt for ecologists. Divers Distrib. 2011;17: 43–57. Available: https://onlinelibrary.wiley.com/doi/abs/10.1111/j.1472-4642.2010.00725.x%4010.1111/%28ISSN%291472-4642.species-distribution-models-in-conservation-biogeography

44. Xu ZL, Ali Z, Yi JX, He XL, Zhang DY, Yu GH, et al. Expressed sequence tag-simple sequence repeat-based molecular variance in two Salicornia (Amaranthaceae) populations. Genet Mol Res. 2011;10: 1262–1276. Available: https://www.geneticsmr.com/sites/default/files/articles/year2011/vol10-2/pdf/gmr1321.pdf

45. Thakur P, Kumar S, Malik JA, Berger JD, Nayyar H. Cold stress effects on reproductive development in grain crops: An overview. Environ Exp Bot. 2010;67: 429–443. doi:10.1016/j.envexpbot.2009.09.004

46. Valentini A, Pompanon F, Taberlet P. DNA barcoding for ecologists. Trends Ecol Evol. 2009;24: 110–117. doi:10.1016/j.tree.2008.09.011

47. Ennos RA. Estimating the relative rates of pollen and seed migration among plant populations. Heredity (Edinb). 1994;72: 250–259. doi:10.1038/hdy.1994.35

48. Slatkin M. ene flow and the geographic structure of natural populations. Science (80-). 1987;236: 787–792.

49. Riccioli F, Fratini R, Boncinelli F, El Asmar T, El Asmar J-P, Casini L. Spatial analysis of selected biodiversity features in protected areas: a case study in Tuscany region. Land use policy. 2016;57: 540–554. doi:10.1016/J.LANDUSEPOL.2016.06.023

50. Vermeulen SJ, Campbell BM, Ingram JSI. Climate Change and Food Systems. Annu Rev Environ Resour. 2012;37: 195–222. doi:10.1146/annurev-environ-020411-130608

51. Kearney M, Porter W. Mechanistic niche modelling: combining physiological and spatial data to predict species’ ranges. Ecol Lett. 2009;12: 334–350. doi:10.1111/j.1461-0248.2008.01277.x

52. Esquinas-Alcázar J. Protecting crop genetic diversity for food security: Political, ethical and technical challenges. Nat Rev Genet. 2005;6: 946–953. doi:10.1038/nrg1729

